# Improving fMRI in Parkinson’s Disease by Accounting for Brain Region-Specific Activity Patterns

**DOI:** 10.1101/2022.08.30.505786

**Authors:** Renzo Torrecuso, Karsten Mueller, Štefan Holiga, Tomáš Sieger, Josef Vymazal, Filip Růžička, Jan Roth, Evzen Růžička, Matthias L. Schroeter, Robert Jech, Harald E. Möller

## Abstract

In functional magnetic imaging (fMRI) in Parkinson’s disease (PD), a paradigm consisting of blocks of finger tapping and rest along with a corresponding general linear model (GLM) is often used to assess motor activity. However, this method has three limitations: *(i)* Due to the strong magnetic field and the confined environment of the cylindrical bore, it is troublesome to accurately monitor motor output and, therefore, variability in the performed movement is typically ignored. *(ii)* Given the loss of dopaminergic neurons and ongoing compensatory brain mechanisms, motor control is abnormal in PD. Therefore, modeling of patients’ tapping with a constant amplitude (using a boxcar function) and the expected Parkinsonian motor output are prone to mismatch. *(iii)* The motor loop involves structures with distinct hemodynamic responses, for which only one type of modeling (e.g., modeling the whole block of finger tapping) may not suffice to capture these structure’s temporal activation. The first two limitations call for considering results from online recordings of the real motor output that may lead to significant sensitivity improvements. This was shown in previous work using a non-magnetic glove to capture details of the patients’ finger movements in a so-called *kinematic approach*. For the third limitation, modeling motion initiation instead of the whole tapping block has been suggested to account for different temporal activation signatures of the motor loop’s structures. In the present study we propose improvements to the GLM as a tool to study motor disorders. For this, we test the robustness of the kinematic approach in an expanded cohort (*n*=31), apply more conservative statistics than in previous work, and evaluate the benefits of an event-related model function. Our findings suggest that the integration of the kinematic approach offers a general improvement in detecting activations in subcortical structures, such as the basal ganglia. Additionally, modeling motion initiation using an event-related design yielded superior performance in capturing medication-related effects in the putamen. Our results may guide adaptations in analysis strategies for functional motor studies related to PD and also in more general applications.

## 1. Introduction

In Parkinson’s disease (PD), the progressive loss of dopaminergic neurons in the substantia nigra typically initiates years before the patient recognizes motor symptoms (Kalia and Lang, 2015) and functioning of the cortico-striato-thalamo-cortical loop (CSTCL) is compromised. Diagnosis may occur many years after the onset of neuronal depletion and, at the time of clinical detection, typically 30% of dopaminergic neurons in the substantia nigra are already lost together with 50– 60% of striatal dopaminergic terminals (Burke and O’Malley, 2013). This ‘silent’ change of the CSTCL leads to brain compensatory mechanisms, such as hyperactivity of the globus pallidus; overactivation of the right dorsolateral prefrontal cortex, the premotor cortex and the supplementary motor area; and excitability increase of the motor cortex (Caproni et al., 2013). Given these alterations and adaptations, functioning of the CSTCL and the patients’ motor output are expected to be unpredictable and to deviate from normal brain motor mechanisms (Albin et al., 1989; Alexander, et al. 1986; DeLong and Wichmann, 2007; Yelnik, 2002).

This unpredictability imposes a methodological obstacle for studies performing functional magnetic resonance imaging (fMRI) in PD patients using finger-tapping paradigms. A considerable number of these studies make use of a general linear model (GLM) with a simple rectangular predictor, referred to as “boxcar function”, which compares measurements from blocks where patients perform a motor task with alternating blocks of rest. The limitation of this predictor, when applied to PD studies, results from the fact that the boxcar function assumes a sustained activity consisting of evenly distributed events of constant amplitude. Clearly, this model has various limitations due to the irregular motor performance within and between blocks of finger tapping. Previously, Holiga et al. (2012) proposed an approach to address the limitation of a heterogeneous motor output by integrating non-magnetic sensory gloves as part of the fMRI exam. The gloves’ fiber-optic sensors collect synchronized, fine-grained movement information from all finger joints, which can be introduced into the GLM design matrix as a more sophisticated regressor. This achieved an increased sensitivity in experiments in a small cohort of 12 patients with PD targeted at the levodopa-related activity in subcortical structures.

In healthy subjects, finger tapping has been related to the activation of putamen (Gatti et al., 2017; Mattay et al., 1998); putamen and caudate nucleus (Bednark et al., 2015; Lewis et al., 2007); or putamen, caudate nucleus and globus pallidus (Lehéricy et al., 2006; Moritz et al., 2000; Riecker et al., 2003). Studies in patients with PD suggest that the ventrolateral layer of the substantia nigra pars compacta, which projects to the putamen’s dorsal and posterior parts, is the most vulnerable circuit to the pathological process. Therefore, the overriding motor deficits in PD have been attributed to the disruption of putamen function (Fearnley and Lees, 1991; Kish et al., 1985; Torstenson et al., 1997). Studies employing motor tasks have also observed higher activity of putamen, caudate nucleus, globus pallidus, and thalamus when comparing patients with PD on and off levodopa medication (Athauda et al., 2017; Brundin et al., 2000; Freeman et al., 1995; Jenkins et al., 1992; Playford et al., 1992; Wu et al., 2012; Yan et al., 2015).

While these studies underline the particular importance of subcortical structures in PD, previous work also indicated further potential methodological limitations. Activity of the putamen in healthy subjects was better captured when finger tapping was modeled as a brief, 2s-long transient function rather than a boxcar function comprising the entire tapping block of tens of seconds (Moritz et al., 2000). This suggests that an extended boxcar function may not capture distinct features of the hemodynamic response function (HRF) among all structures of the CSTCL.

Following up on Holiga et al.’s approach, we conducted fMRI measurements in PD patients with a finger-tapping task in a levodopa-on and off state (further denoted by “i-DOPA ON” and “l-DOPA OFF”, respectively), in which motor performance was simultaneously assessed by a non-magnetic sensory glove. After converging the time series of sensor readings into a predicting regressor, we compared it with a constant-amplitude boxcar function convolved with the canonical HRF without integration of the information obtained with the glove. Beyond a verification of earlier results (Holiga et al., 2012), our goals were to examine if an expanded cohort (*n*=31) might allow an identification of further aspects of the motor system in PD and, at the same time, to employ more conservative statistics for improved robustness of the findings. Finally, we performed analyses employing a so-called *event-related* (ER) design that considered the onsets of tapping and rest comparing these results to both the standard block-design approach based on a boxcar function and the kinematic model based on the glove readings. We hypothesized that different models may be more suitable for detecting activation in specific brain regions and l-DOPA drug effect.

## 2. Material and Methods

### 2.1 Subjects

The study had been approved by the Ethics Committee of the General University Hospital in Prague, Czech Republic, in accordance with the Declaration of Helsinki. A total of *n*=31 right-handed patients with PD (26 males, age 56.8±7.7 years) of equivalent akinetic-rigid type, Hoehn-Yahr stages II-III (Hoehn and Yahr, 1967) were included in our study after giving informed written consent. This “combined cohort” included a subset of *n*=11 patients (subsequently referred to as “initial cohort”) from a previously published investigation (Holiga et al., 2012), as well as *n*=20 newly recruited patients (subsequently referred to as “new cohort”). Note that the data from one patient of Holiga et al.’s cohort (12 patients) could not be integrated because of a corrupted file. The patients’ motor symptom severity was clinically assessed using the Unified Parkinson’s Disease Rating Scale III (UPDRS-III) (Ramaker et al., 2002). Disease duration was 12.3±2.5 years, and levodopa treatment duration was 9.2±2.9 years. More characteristics of the combined cohort are summarized in Table 1.

**Table 1.**
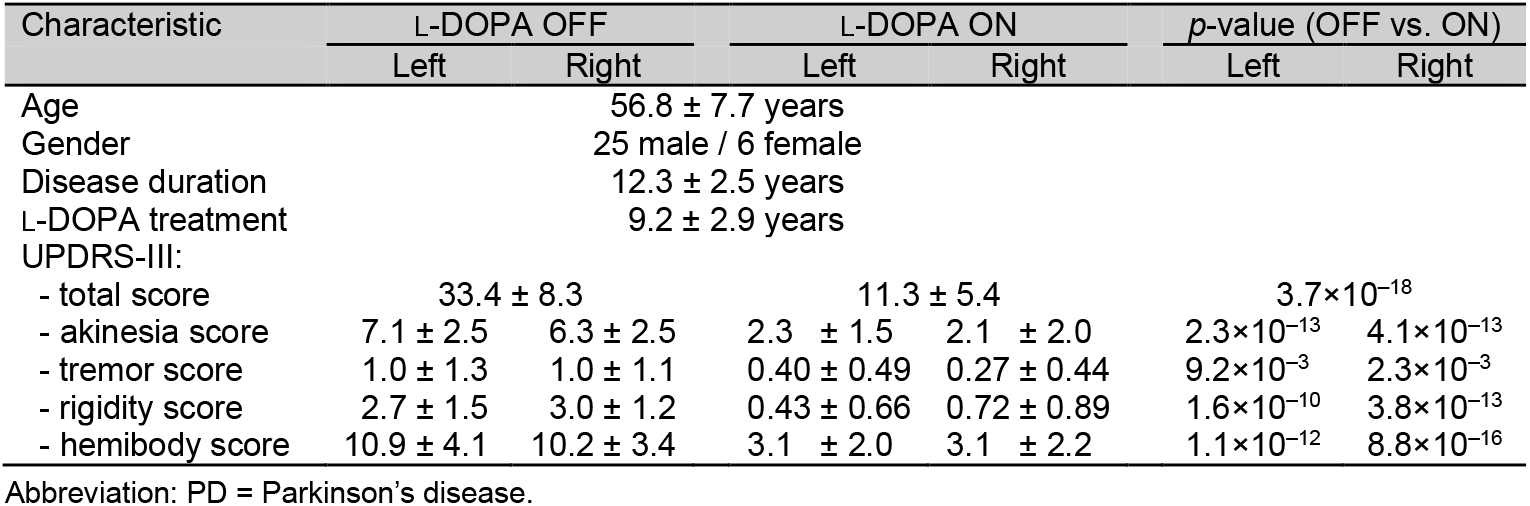
Clinical characteristics (mean values plus/minus one standard deviation) of the combined cohort of *n*=31 PD patients included in the study.

### 2.2 Experimental Design and Data Acquisition

All fMRI exams were performed on a 1.5T MAGNETOM Symphony scanner (Siemens Healthineers, Erlangen, Germany) using a birdcage head coil and a gradient-echo echo planar imaging (EPI) sequence (Mansfield, 1977) with a repetition time of *TR* = 1 s, an echo time of 54 ms, and a flip angle of 90°. Ten coronal slices (thickness 3 mm, gap 1 mm, nominal in-plane resolution 3×3 mm^2^) were obtained covering the basal ganglia and the primary motor cortex (M1).

All patients performed a task that consisted of 25 tapping periods of 10 s each alternating with 25 rest periods of the same duration, yielding a total duration of 500 s per session. Both hands were investigated in separate runs using the same paradigm. In order to support tapping regularity, visual cues blinking at 1 Hz were delivered to the patients via a projection screen. Each patient performed both runs of the entire task twice, during two separate scanning sessions. The first session was performed after a one-night withdrawal of levodopa intake, and the second session occurred one hour after administration of 250 mg levodopa/25 mg carbidopa (Isicom 250, Desitin Arzneimittel, Hamburg, Germany).

While performing the task inside the scanner, the patients used a non-magnetic glove containing 14 fiber-optic sensors with 64Hz sampling rate (5th Dimension Technologies, Irvine, CA, USA). The sensors captured adduction and abduction within neighbor fingers as well as individual flexion and extension of each finger. The patients were instructed to tap the index finger of one hand in opposition to the thumb. The glove regressor that was used as an input to the individual-level fMRI design matrix, was computed by averaging each session’s time course from the 14 glove sensors into a single waveform. Before fMRI scanning, a normalization procedure was conducted, in which all patients were requested to perform calibration gestures to establish their individual peak amplitude and baseline values.

### 2.3 Medication Effect on UPDRS-III Scores and Kinematics

An initial evaluation of the variability of the simultaneous recordings from the full set of sensors indicated that all sensors captured the tapping movement. The sensor on the index finger’s proximal interphalangeal joint delivered non-redundant movement information with the highest signal amplitude, and was used for a behavioral analysis of a potential l-DOPA effect. In particular, we established a comparison between the patients’ UPDRS-III akinesia ratings off and on medication with further parameters characterizing the tapping performance. In order to investigate a potential l-DOPA effect, *(i)* the variance of the amplitude within each block and *(ii)* the reaction time between the visual cue and the first tap were computed and averaged over all blocks (Supplementary Figure S1):

### 2.4 fMRI Data Analysis

All fMRI data analyses were performed using SPM12 (www.fil.ion.ucl.ac.uk/spm/) with Matlab R2017b (MathWorks, Natick, MA, USA). Image pre-processing included realignment for motion correction, normalization to the MNI (Montreal Neurological Institute) space using a separately acquired individual *T*_1_-weighted MP-RAGE (Magnetization-Prepared RApid Gradient Echo) (Mugler and Brookeman, 1990) dataset, and smoothing with a Gaussian spatial filter of 10mm full width at half maximum (FWHM).

For evaluating the impact from integrating the glove information into the analysis pipeline, the data were processed twice: *(i)* using a GLM without consideration of the glove information using a boxcar function (“standard model”), and *(ii)* using a GLM with the glove information as a regressor instead of a static boxcar function (“glove model”). The standard model consisted of the conventional SPM’s first-level block design employing a boxcar function according to the specifications of the finger-tapping blocks’ onsets and durations (10 s for both tapping and rest blocks), convolved with the canonical HRF. The design function for the glove model was based on the glove sensors’ readings of individual taps convolved with the canonical HRF. Compared to previous work (Holiga et al. 2012), the processing pipeline was slightly modified to include the following adaptations that better reflect current standards in fMRI analysis: *(i)* use of SPM12 instead of SPM8, *(ii)* consideration of six head-motion regressors in the GLM design matrix in all analyses, *(iii)* use of a smoothing kernel with 10mm FWHM instead of 8 mm, and *(iv)* resampling to 2×2×2 mm^3^ instead of 3×3×3 mm^3^ after normalization (Mueller et al., 2017).

For disentangling activity due to the *initiation* of the tapping as compared to the *sustained execution* of the entire series of taps during each block, further analyses were performed employing an ER approach that was entirely based on the onset of tapping and rest epochs. In particular, a conventional SPM’s first-level design was obtained by the specifications of the finger tapping blocks’ onsets, however, with the block durations set to zero. In this case, instead of modeling tapping and rest in a same predictor vector, the SPM’s first-level analysis was modeled using a contrast between two vectors, one reflecting the tapping blocks onsets minus another one reflecting the rest block onsets. Subsequently, we refer to this approach as “onset model” in order to distinguish it from the glove model that considers a series of events in relatively rapid succession.

After parameter estimation, *β*-images were obtained for a second-level statistical analysis. One-sample *t*-tests were conducted across the *β*-images of the l-DOPA ON and OFF conditions for both hands (left and right) separately. In order to assess group-level activation maps showing the l-DOPA effect, a flexible factorial analysis (FFA) was carried out using a two-by-two design (l-DOPA ON/l-DOPA OFF; left hand/right hand) as main effect of both factors. Thereafter, contrast images were generated from the *β*-images investigating potential differences between l-DOPA ON and l-DOPA OFF.

All second-level analyses (one-sample *t*-tests as well as the FFA) were performed for the standard and glove model as well as for the onset model. For a direct comparison with previous results, the initial cohort (*n*=11) and the new cohort (*n*=20) were analyzed separately applying the same significance threshold (*p*<0.001, uncorrected) as used by Holiga et al. (2012). This procedure was lately referred to as *‘cluster defining threshold’* (Eklund et al., 2016). In addition to this cluster-wise inference, we further applied a voxel-wise inference, in which only results were regarded as significant with an error probability of *p*<0.05 after family-wise error (FWE) correction at the *voxel level* (Mueller et al., 2020). All analyses were performed with cluster extent set to 30 voxels.

In order to investigate the temporal activation signatures of different CSTCL structures, we further performed an additional finite impulse response (FIR) analysis. Instead of modeling an entire tapping block as a boxcar function of corresponding duration, the FIR models the BOLD signal by applying contiguous boxcar functions of much shorter duration, typically set to one *TR*. This procedure allows to average the signal as a parameter estimate at each *TR* and, thereby, to compare how different regions of interest (ROIs) perform in time (Friston, 2003; Henson et al., 2001; Nakamura et al., 2012).

## 3. Results

### 3.1 Medication Effect on Clinical Scores

The clinical assessments revealed l-DOPA effects on all UPDRS-III (sub)scores corresponding to significantly reduced symptoms in the l-DOPA ON condition (Table 1 and Figure 1). The comparisons of the behavioral data (session average values of the tapping characteristics) revealed an l-DOPA effect for the left hand’s average amplitude variance, whereas there was no significant response-time difference of the two l-DOPA conditions (Table 2). A comparison of the two cohorts (two-sample *t*-test, “initial” vs. “new”) revealed insignificant differences in all parameters except of a disparity in the tremor scores of the right hand, which was lower in the initial cohort (l-DOPA OFF: 0.6±0.7 vs. 1.6±1.4, *p*=0.01; l-DOPA ON: 0.15±0.3 vs. 0.5±0.5, *p*=0.03). Given the generally low tremor scores in all patients, we do not expect a relevant impact from this subtle difference.

**Figure 1.**
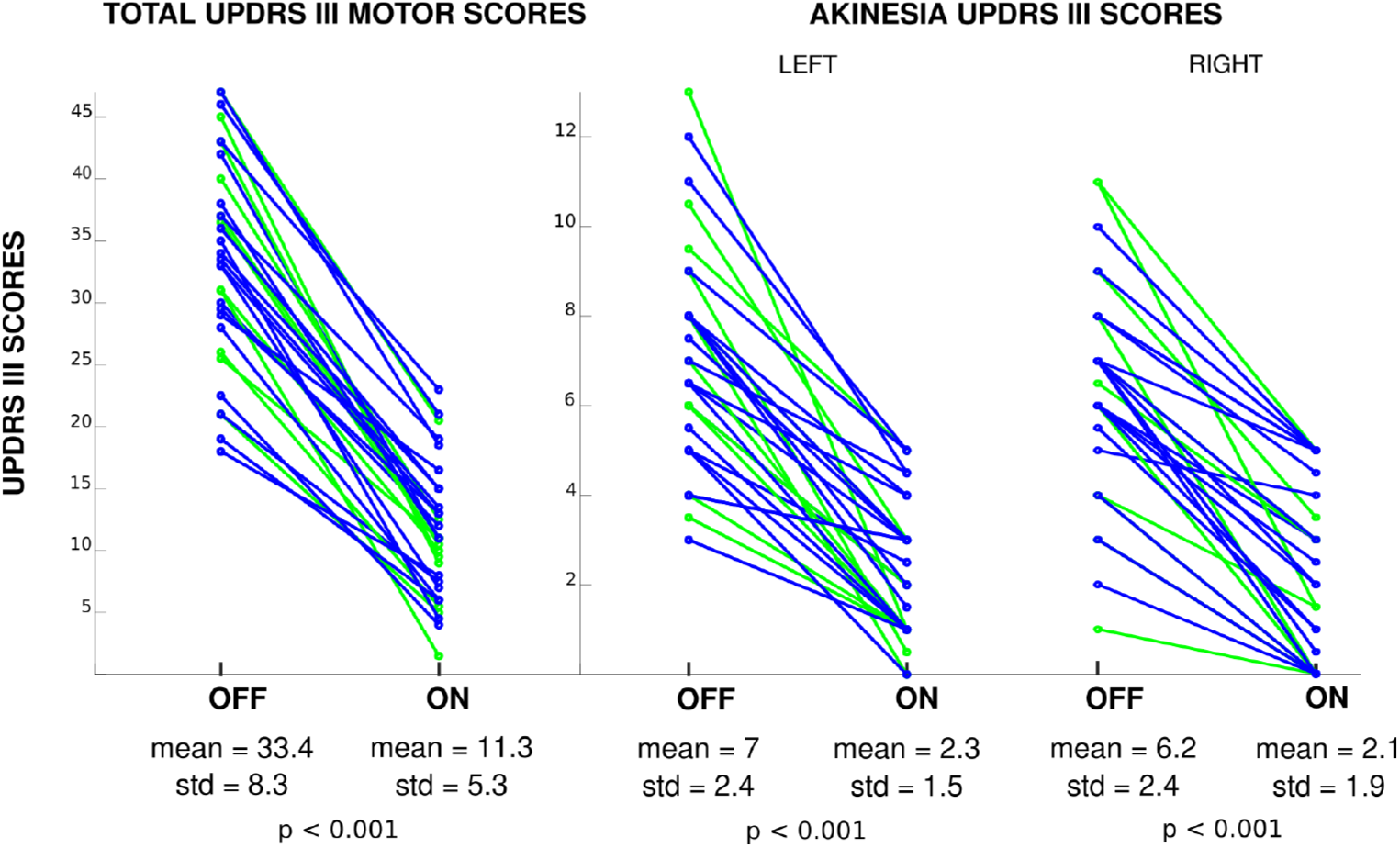
UPDRS-III rating results including the total motor scores and the akinesia subscores of the left and the right side in all patients of the combined cohort comparing the l-DOPA OFF and l-DOPA ON conditions.

**Table 2.** Group-level results (combined cohort, *n*=31) of averaged tapping parameters (mean plus/minus one standard deviation) recorded with the sensor on the index finger’s proximal interphalangeal joint (left and right hand) during the “l-DOPA OFF” and “l-DOPA ON” conditions.

| Parameter | Hand | L-DOPA OFF | L-DOPA ON | $p$-value |
| --- | --- | --- | --- | --- |
| Amplitude variance<sup>a</sup> / a.u. | Left | 200 ± 200 | 670 ± 1000 | **0.01** |
|  | Right | 750 ± 950 | 1300 ± 1700 | 0.12 |
| Response delay / ms | Left | 10.5 ± 8.5 | 11.2 ± 6.2 | 0.71 |
|  | Right | 10.5 ± 11.9 | 11.0 ± 5.7 | 0.82 |

<sup>a</sup> Average amplitude variance within a block. Abbreviation: a.u. = arbitrary units.

### 3.2 fMRI Main Effects of Tapping versus Rest

The analysis of the fMRI data showed significant brain activity differences related to finger movement: For finger tapping with the (dominant) right hand in the l-DOPA ON condition, a group-level one-sample *t*-test in the initial cohort with the standard block-design model revealed a cluster of activity in the primary cortical motor regions (Figure 2a, and Table 3). The same result was also obtained with the glove model, however, with substantially increased cluster size (approx. sixfold) and peak *t*-values underlining a marked gain in sensitivity for detecting activation in cortical motor regions achieved with kinematic modeling. This verification of Holiga et al.’s observation in the same cohort was reproduced with the new cohort with an even larger gain in cluster size for the glove compared to the standard model (Figure 2d,e), in line with an increased statistical power achieved with the larger cohort (*n*=20 vs. *n*=11). Consistently, a major sensitivity increase was obtained in the combined cohort (*n*= 31; Figure 2g,h). Further results obtained with the increased sample size of the combined cohort comprised activations in subcortical areas of the motor loop, including the left putamen and thalamus and the right caudate nucleus. Note that these subcortical findings were only significant when employing the glove model but were not obtained with the standard model (Table 3).

**Figure 2.**
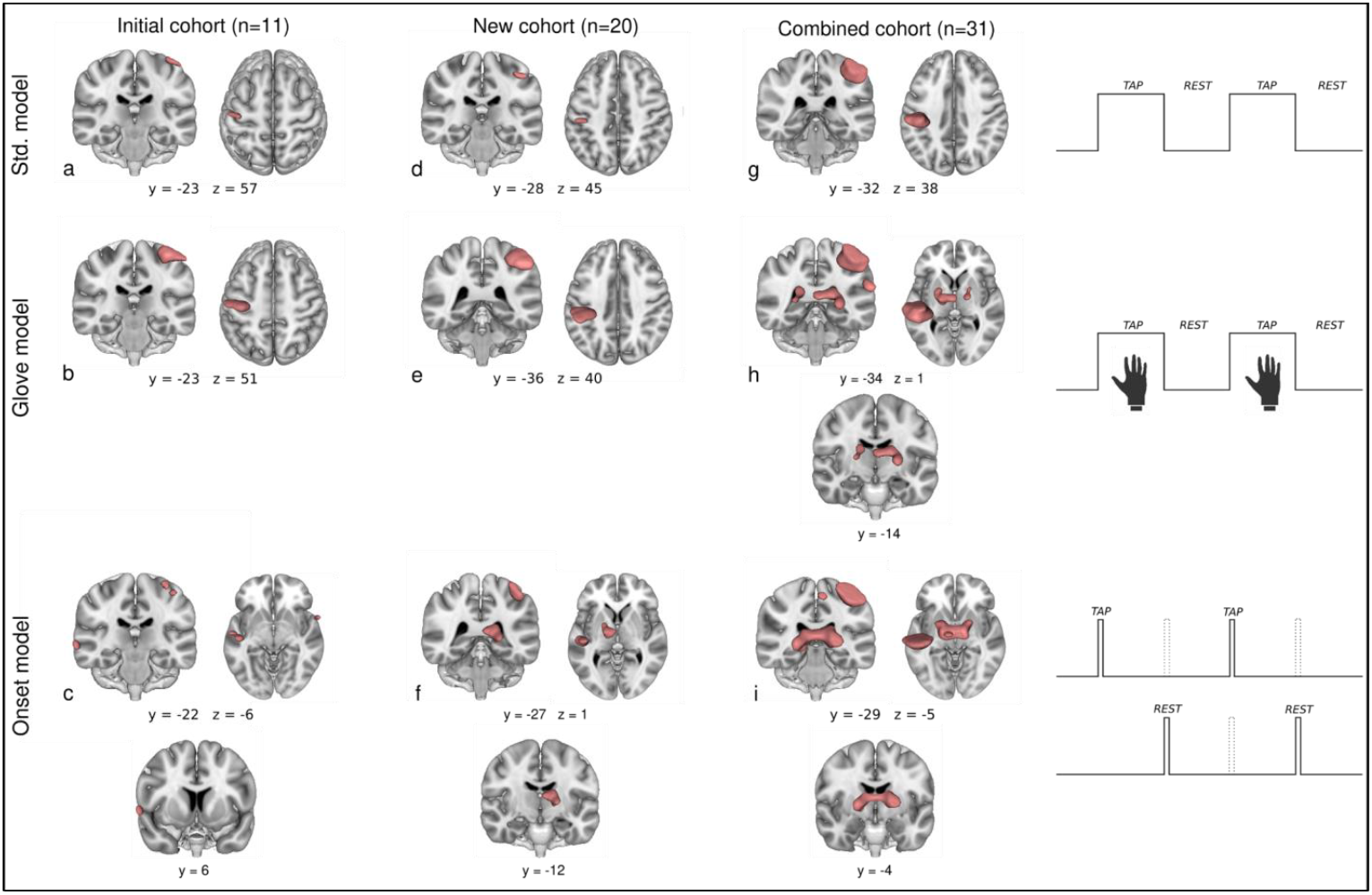
Activation maps (one sample *t*-test, main effect of tapping vs. rest; group results) for finger tapping with the (dominant) **right hand** (l-DOPA ON) obtained in the initial (*n*=11) **(a–c)**, new (*n*=20) **(d–f)**and combined cohort (*n*=31) **(g–i)**with the standard model (block design employing a boxcar function) **(a, d, g)**, the glove model **(b, e, h)**, and the onset model **(c, f, i)**. The coordinates refer to the displayed anatomical slices and not to clusters’ maxima. Further quantitative results are summarized in Table 3.

**Table 3.**
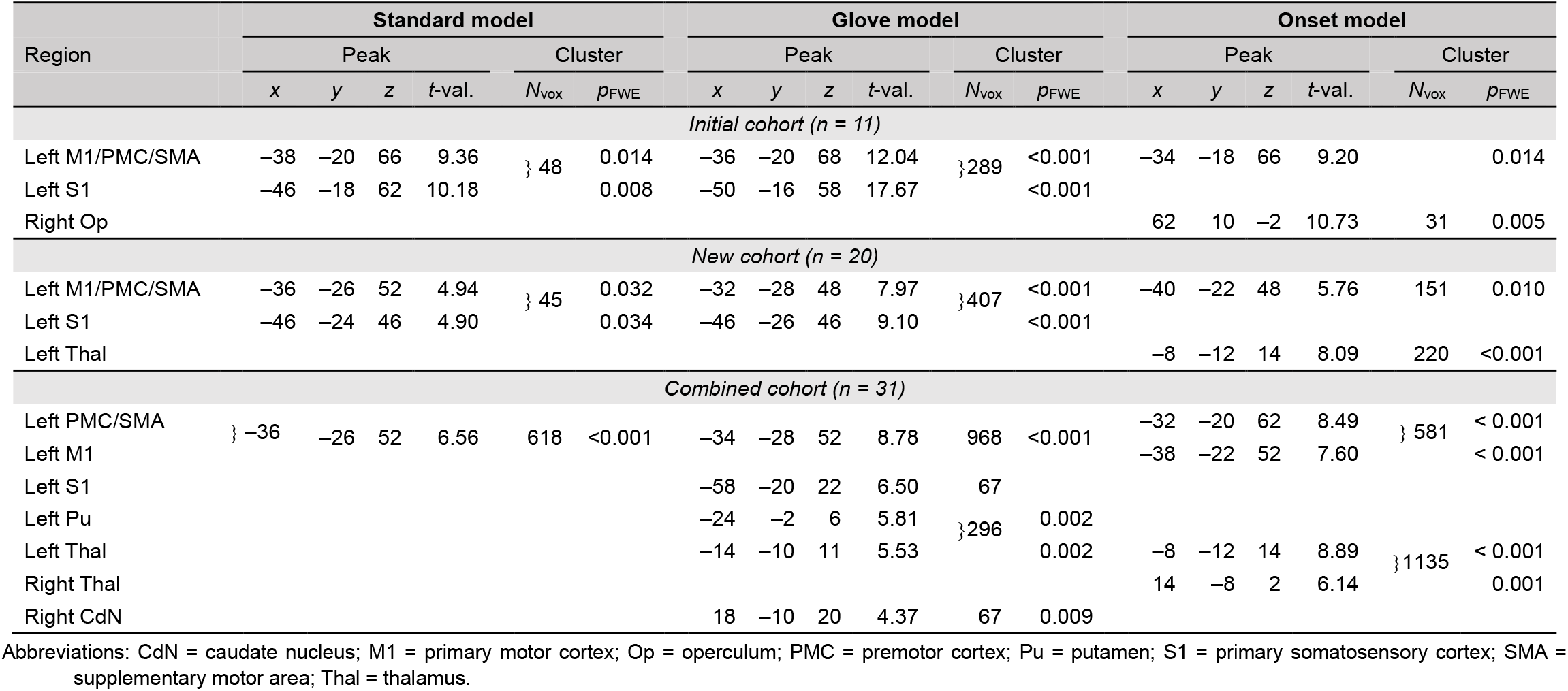
One-sample *t*-test results (MNI coordinates in mm and *t*-values of the peak activation in a cluster as well as cluster sizes and FWE-corrected *p*-values) for finger tapping with the (dominant) **right hand** (l-DOPA ON condition) obtained in the initial (*n*=11), new (*n*=20) and combined cohort (*n*=31) with the standard model (block design employing a boxcar function), the glove model, and the onset model. Corresponding activation maps are shown in Figure 2.

Compared to both the standard and the glove model, the onset model detected also substantial subcortical activity, yielding a cluster in the left thalamus already with the new cohort and a substantially increased cluster comprising bilateral thalamus regions with the larger combined cohort (Figure 2f,i and Table 3). Remarkably, the opposite trend was obtained in cortical regions, that is, activations in M1 identified with the onset model were markedly reduced as compared to the glove and the standard model results. No *t*-value improvement was achieved upon integrating the glove data into the onset model.

For tapping with the (non-dominant) left hand (l-DOPA ON condition), similar results (i.e., contralateral activation in motor areas) as for right-handed tapping were obtained, however, with lower *t*-values and overall smaller cluster sizes (Supplementary Figure S2 and Supplementary Table S1). In this case, significance was not reached in subcortical areas, including the combined cohort.

Further insight was obtained from the FIR analysis showing that putamen activation peaked around the onset of tapping (Figure 3). Upon graphically overlaying this result with SPM’s tapping predictor, it is evident that the onset model achieves better fits of the temporal activation signature in the putamen than the block design (Figure 3c). Note that the opposite behavior was evident for activity in M1 (Figure 3d), that is a sustained activity throughout the tapping block that is best captured by the block design.

**Figure 3.**
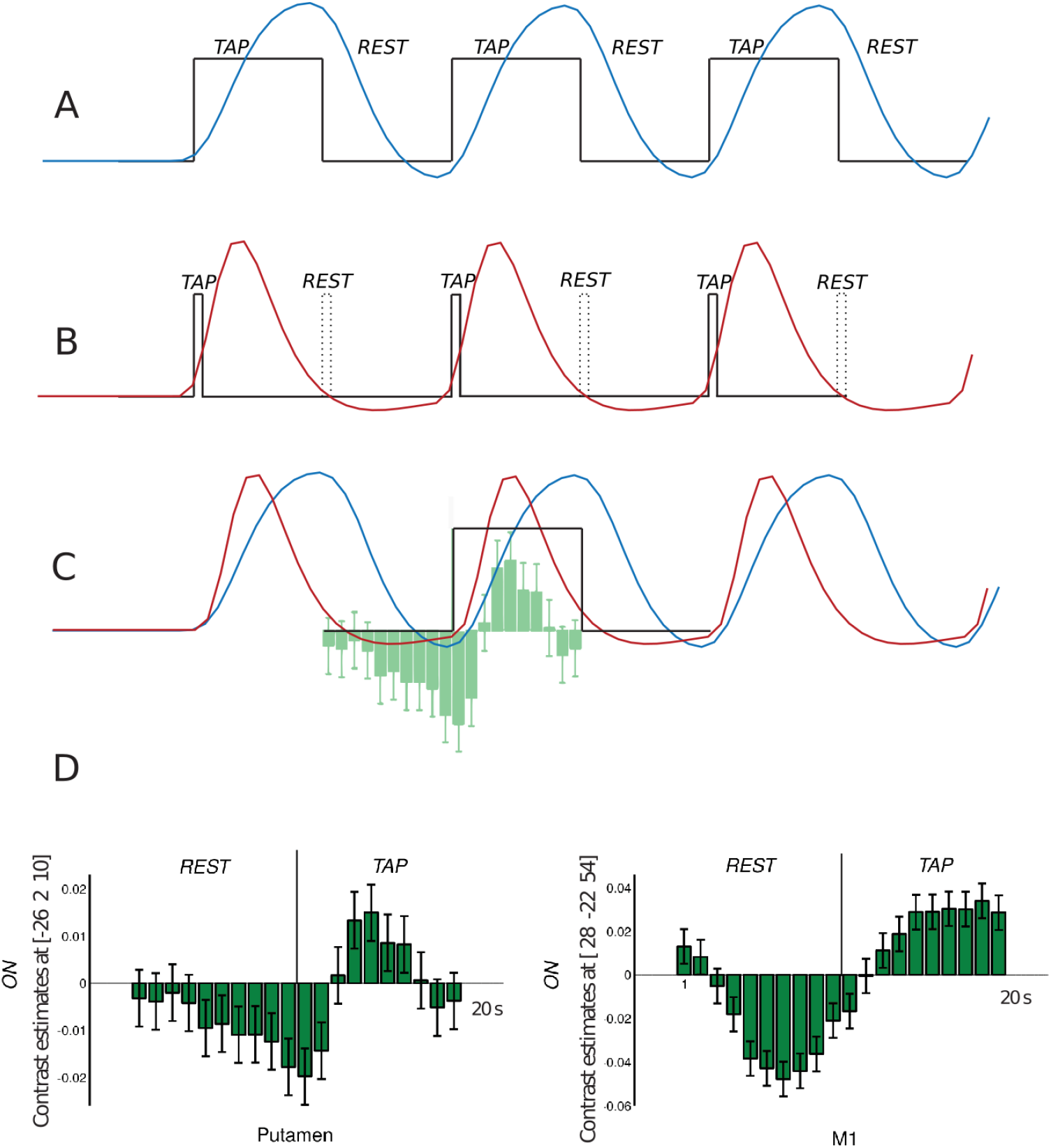
Depiction of model functions used in block and ER designs. **(A)** Following convolution with the HRF, the block design leads to a model function (blue solid line) that reaches its peak around the end of the 10s-tapping block, whereas **(B)** a design considering only a brief event at tapping onset (black solid line) produces a corresponding model function (red solid line) that peaks approximately 5 s after tapping onset. Note that the blue and red curves were directly extracted from SPM’s first-level computation employing the characteristic timing of the paradigm (amplitudes are adjusted for better visualization). **(C)** An overlay of both design functions and the results from a FIR analysis demonstrates that an onset model based on an ER design is better suited to capture activity in the putamen. **(D)** Comparison of the FIR analysis results obtained in the putamen **(left)** and M1 **(right)**. While M1 presents a more sustained activation throughout the block, putamen activity peaks around 5 s followed immediately by a decay.

### 3.3 Effect of l-DOPA Medication on fMRI Results

The FFA computing a contrast between the l-DOPA ON and the l-DOPA OFF treatment state yielded distinct differences in the activation maps obtained with the three models, with higher activity observed for l-DOPA ON (Figure 4 and Table 4): *(i)* With the standard model, no result survived FWE correction (*p*<0.05 at the voxel level) in any of the cohorts, yielding an “empty brain”. *(ii)* The glove model applied to the combined cohort (*n*=31) revealed activation of the left and right putamen, with the cluster in the left putamen extending into the left thalamus. For the smaller initial (*n*=11) and the new cohort (*n*=20), however, no activation survived FWE correction. *(iii)* Prominent activations centered on the left and right putamen were obtained in all cohorts with the onset model, with increasing *t*-values and cluster extents obtained upon increasing the sample size (Figure 4).

**Figure 4.**
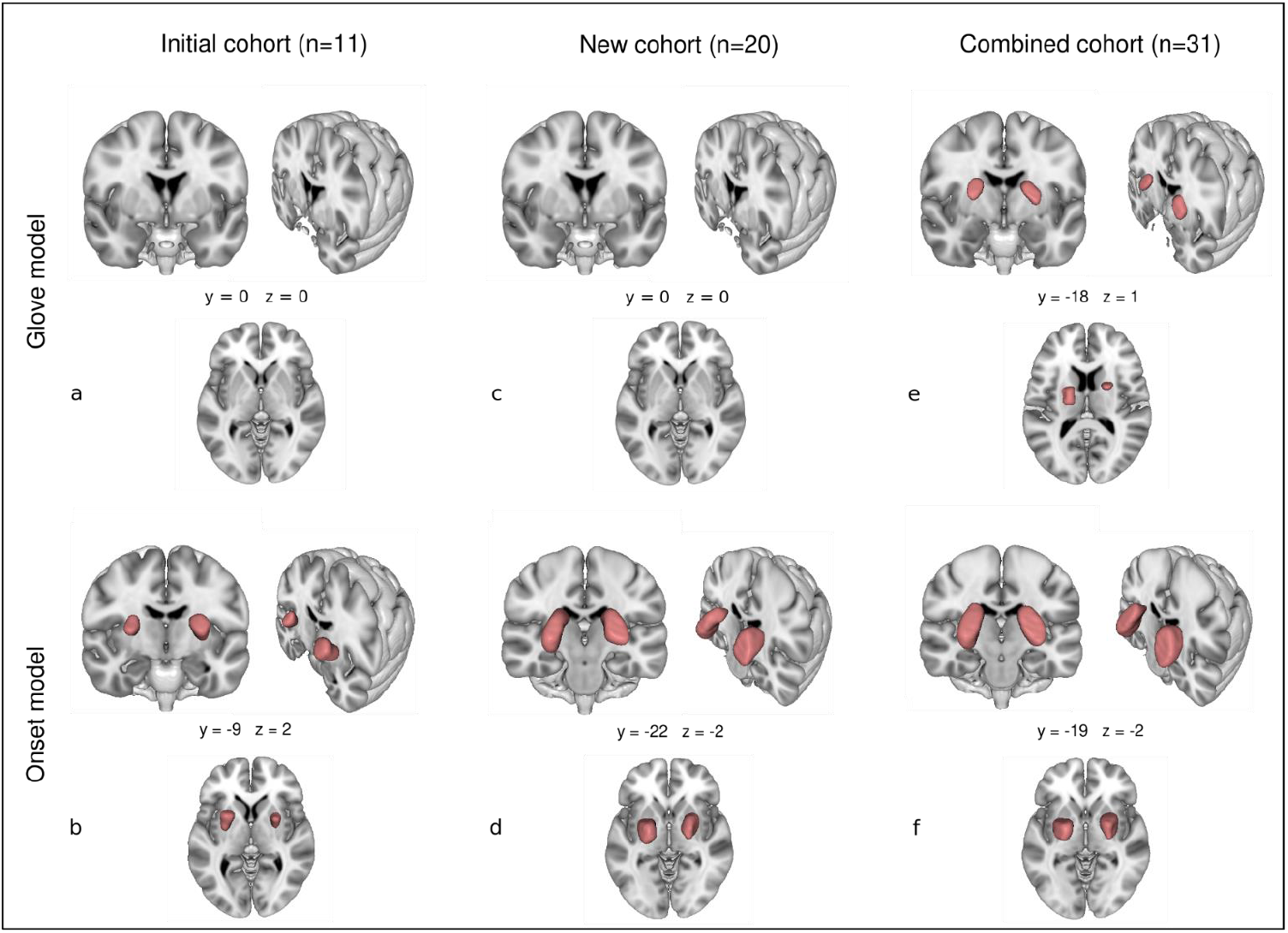
Activation maps (FFA, main effect of l-DOPA ON vs. OFF) for finger tapping obtained in the initial (*n*=11) **(a, b)**, new (*n*=20) **(c, d)** and combined cohort (*n*=31) **(e, f)** with the glove model **(a, c, e)** and the onset model **(b, d, f)**. With the standard model (block design employing a boxcar function), no activation survived the FWE correction (results not shown here). The coordinates refer to the displayed anatomical slices and not to the clusters’ maxima. Further quantitative results are summarized in Table 4.

**Table 4.** Flexible factorial analysis results (l-DOPA ON vs. OFF condition) for finger tapping (MNI coordinates in mm and *t*-values of the peak activation in a cluster as well as cluster sizes and FWE-corrected *p*-values) obtained in the initial (*n*=11), new (*n*=20) and combined cohort (*n*=31) with the glove model and with the onset model. Corresponding activation maps are shown in Figure 4. Note that no activations were detected with the standard model.

| Region | Glove model |  |  |  |  |  | Onset model |  |  |  |  |  |
| --- | --- | --- | --- | --- | --- | --- | --- | --- | --- | --- | --- | --- |
|  | Peak |  |  |  | Cluster |  | Peak |  |  |  | Cluster |  |
| | x | y | z | t-val. | $N_{\text{vox}}$ | $p_{\text{FWE}}$ | x | y | z | t-val. | $N_{\text{vox}}$ | $p_{\text{FWE}}$ |
| *Initial cohort (n = 11)* |  |  |  |  |  |  |  |  |  |  |  |  |
| Left Pu |  |  |  |  |  |  | -24 | 8 | 14 | 6.65 | 291 | <0.001 |
| Right Pu |  |  |  |  |  |  | 26 | 4 | 12 | 5.96 | 135 | 0.002 |
| *New cohort (n = 20)* |  |  |  |  |  |  |  |  |  |  |  |  |
| Left Pu |  |  |  |  |  |  | -22 | -2 | 8 | 6.53 | 633 | <0.001 |
| Right Pu |  |  |  |  |  |  | 24 | 2 | 4 | 6.88 | 629 | <0.001 |
| *Combined cohort (n = 31)* |  |  |  |  |  |  |  |  |  |  |  |  |
| Left Pu | -18 | 2 | 10 | 4.96 | } 232 | 0.002 | -24 | -2 | 10 | 8.13 | 665 | <0.001 |
| Left Thal | -20 | -16 | 14 | 4.92 |  | 0.003 |  |  |  |  |  |  |
| Right Pu | 22 | 4 | 16 | 5.33 | 91 | 0.001 | 26 | 2 | 6 | 7.94 | 790 | <0.001 |

Abbreviations: Pu = putamen; Thal = thalamus.

## 4. Discussion

This study was conceived as an investigation of potential benefits from fine-tuning the brain signal modeling and analysis strategy in fMRI studies of patients with PD performing a simple finger-tapping task. For this purpose, we focused on *(i)* an evaluation of the usefulness of kinematic modeling utilizing information from parallel recordings of finger movements with sensory gloves and *(ii)* an adaption of the standard design function to particularly focus on the onsets of tapping and rest blocks. To assess the robustness of such observations in relatively small cohorts that are typically available in clinical applications of fMRI, we re-analyzed data from previous experiments (“initial cohort”, *n*=11) (Holiga et al., 2012) applying current state-of-the art statistics and compared the results to those from a “new cohort” (*n*=20) as well as a larger “combined cohort” (*n*=31).

The comparison of the tapping performance (i.e., glove recordings) for the l-DOPA OFF and l-DOPA ON conditions in the combined cohort revealed a significant difference in the tapping amplitude variance, specifically, in the non-dominant hand. This may be due to long-term motor compensation processes parallel to the continuous degeneration of nigral-putamic dopamine supply. With the latter’s restoration in the l-DOPA ON condition, the left hand’s movement is unleashed but, as its CSTCL has not received a comparable degree of daily movement adaptation as for the dominant hand, tapping is expected to perform in a less controlled way.

To the best of our knowledge, no further study has evaluated kinematic modeling in the context of fMRI studies in patients with PD since the initial publication by Holiga et al. (2012). Three recent meta-analyses (Herz et al., 2014, 2021; Spay et al., 2019) summarize that finger-tapping protocols in the context of PD compared patients on and off levodopa treatment or patients and healthy controls, all without considering potential limitations due to the application of a standard block design. This approach employs a boxcar function to capture the BOLD signal from a sustained repetitive task, intercalated with resting blocks, when the modeling function’s amplitude returns to baseline (Macey et al., 2016). Inherently, this model predicts that the tapping is performed regularly, which is a meaningful approximation for healthy subjects. However, patients with PD are typically characterized by an uneven execution of the task as evidenced by the glove data. Consequently, accounting for the real tapping performance by kinematic modeling achieved substantial improvements in the robust detection of activation in areas involved in the execution of movements including M1, premotor cortex and the supplementary motor area as well as sensory input, such as the primary somatosensory cortex (S1). With 14 sensors and a 64 Hz sample rate, the glove provides fine grained spatial-temporal movement information that may be taken as reflecting the ground truth. Integrating this information into the fMRI analysis did not only improve the detection of activation when contrasting tapping versus rest but also the investigation of the medication effect (l-DOPA ON vs. l-DOPA OFF) with the FFA. Consequently, CSTCL structures, such as putamen, expected to emerge in motor paradigms were not observed with the standard model but were clearly identified in the combined cohort when employing the glove model with rather strict statistics (voxel-level FWE correction) that minimizes false-positive observations. This is of particular importance in a scenario where fMRI is employed to evaluate treatment effects as the lack of detecting basal ganglia activation due to an analysis approach that is penalized by degraded sensitivity may be mistaken for the absence of a treatment response.

The results obtained with the onset model suggest that detection of basal ganglia activation is even more sensitive upon restricting the tapping block to its onset while ignoring the sustained repetitions of the tapping events. Regarding activity in putamen, this model outperformed the glove model as evidenced by the fact that robust identification of a drug effect was obtained already with a rather small sample of only 11 patients and further corroborated with larger cohorts. This is in line with earlier observations in healthy subjects by Moritz et al. (2000) demonstrating improved sensitivity in subcortical brain regions when assuming an initial transient response, like our onset model.

The results obtained with the FIR analysis provided means to assess the suitability of each particular model in a specific ROI. While the onset model led to improved statistics in putamen but not in M1, the opposite was true for the glove and the standard model. Furthermore, integration of the glove data did not lead to a relevant improvement of the onset model. This is likely due to the long latency and broadened shape of the HRF, which occur on a timescale of the order of 5 s (Friston, 2003). Therefore, the subtle correction of the occurrence of every block’s first tap (order of 100 ms) that can be obtained from the glove information is smeared away upon convolution with the HRF. Hence, to capture activation in the putamen, which peaks around 5 s after initiation of tapping and then decays back towards the baseline, the glove information provided only minimal benefit. Conversely, for M1, which is characterized by sustained activation throughout the entire tapping block, the glove data permit an individual refinement of the model function for every block. Due to these distinct differences in their activation patterns, activity in the putamen is more appropriately described by the onset model whereas a relevant improvement of the detection of M1 activation is obtained with the glove model.

Previous fMRI literature investigating patients with PD performing motor tasks under different treatment conditions demonstrated deviating findings (see, e.g., Spay et al., 2019 for a comprehensive review), which is probably partly due to variability in the experimental design, selection of ROIs, or cohort sizes. Nevertheless, in view of our current results, we cannot exclude the possibility that activation might have remained undetected due to suboptimal design functions for particular experimental conditions or target regions. In line with the onset-model results (as well as the glove model in a sufficiently large cohort), the putamen appears as a central structure in the restoration of the nigral dopaminergic supply promoted by levodopa, corroborating classical descriptions of the CSTCL (Alexander and Crutcher,1990; Graybiel 1998).

The emergence of activation in the putamen in response to dopaminergic treatment is, hence, a plausible drug effect that is best captured with the onset model. This suggests a role in the preparation for movement rather than in sustained activity related to the execution of repeated tapping events.

Due to the complexity, but also flexibility, of fMRI data-processing tools, there is a debate about the reliability and stability of fMRI results in general. Recently, Botvinik-Nezer et al. (2020) set out to investigated the variability of fMRI results by asking 70 independent research teams to perform an analysis of the same dataset testing the same predefined hypotheses. They found major differences in the individual results leading to substantial impact on the scientific conclusions. Thus, information on the validity and robustness of fMRI results is of paramount interest. Notably, the replication part of the current study adds to this discussion in multiple ways: *(i)* By re-analyzing previous data (Holiga et al., 2012) employing newer software tools and currently suggested parameter settings; *(ii)* by repeating the identical experiment in a new cohort; and *(iii)* by analyzing the merged cohorts with particularly conservative statistics. All three analyses yielded consistent findings, namely an increased activity in the putamen with levodopa administration—albeit with expected differences in the statistical power reflecting the different cohort sizes. Conceptually, we further compared different design functions based on considerations about specific activity patterns in different regions of the motor loop. Again, these adaptations produced consistent results in all cohorts, but also hinted at the importance of adequate models of activity patterns, which may differ between brain regions that are activated in the same experiment. Remarkably, Botvinik-Nezer et al. (2020) obtained a significant consensus in activated regions across teams employing meta-analysis techniques, which goes in line with our results obtained with different analysis pipelines and cohorts. Further work is necessary to clarify the robustness of fMRI in terms of signal strength and signal-to-noise ratio.

Despite the relatively large cohort size (considering application of fMRI in a clinical context) and agreement with the well-established literature on CSTCL functioning in healthy individuals, we recognize the lack of a control group as a limitation of our study. That could have added more robustness to the interpretation of transient activation in specific ROIs and provided further information to which extent patterns of activity in the CSTCL might differ between medicated patients with PD and healthy individuals.

## 5. Conclusion

This study proposes conceptual and methodological improvements of fMRI studies investigating motor performance. In particular, we found that kinematic modeling of finger tapping outperformed the standard approach that models blocks of tapping and rest as a simple convolution of a boxcar function convolved with the HRF. This result obtained with voxel-wise inference corroborates previous findings that had only been achieved with a less robust cluster-defining threshold. Considering the movement initiation impairment that is characteristic for PD, we further applied an analysis strategy that focused on the onset of tapping and rest blocks yielding superior performance in detecting activation of the putamen. Taken together, this suggests that the detection of activation in different brain areas may benefit from different analysis strategies adapted to their particular role. In the current case, the onset model is the preferred choice for capturing the L-DOPA effect on putamen activity, while the glove model yields the best results for detecting M1 activity. Conceptually, these findings advocate careful consideration of individual brain region’s responses in the fMRI analysis strategy.

## Sample CRediT author statement

Renzo Torrecuso: Methodology, software, validation, formal analysis, writing original draft, review & editing, visualization;

Karsten Mueller: Conceptualization, methodology, software, formal analysis, writing original draft, review & editing, supervision, project administration;

Štefan Holiga: Software, review & editing;

Tomáš Sieger: Software, data curation, writing;

Josef Vymazal: Review & editing, resources;

Filip Růžička: Review & editing, resources, data curation;

Jan Roth: Review & editing;

Evzen Růžička: Review & editing;

Matthias L. Schroeter: Funding acquisition, review & editing;

Robert Jech: Conceptualization, resources, data curation, review & editing, project administration, funding acquisition;

Harald E. Möller: Conceptualization, methodology, writing original draft, review & editing, supervision, project administration, funding acquisition.

## Acknowledgements

This work has been supported by a grant of the National Institute for Neurological Research, Czech Republic, Programme EXCELES (ID project No. LX22NPO5107) to RJ, with further funds received from the European Union (NextGenerationEU package) and the Charles University: Cooperatio Program in Neuroscience, and the European Joint Programme Neurodegenerative Disease Research (JPND) 2020 call “Novel imaging and brain stimulation methods and technologies related to Neurodegenerative Diseases”-Neuripides project. MLS was supported by the Deutsche Forschungsgemeinschaft (SCHR 774/5-1) and the eHealthSax Initiative of the Sächsische Aufbaubank (SAB). RT acknowledges support from the International Max Planck Research School on Neuroscience of Communication: Structure, Function, and Plasticity (IMPRS NeuroCom).

