## Supplementary material for "Improving fMRI in Parkinson’s Disease by Accounting for Brain Region-Specific Activity Patterns"

**Abbreviations & Symbols**

BOLD = blood oxygenation level dependent; CdN = caudate nucleus; CSTCL = cortico-striatal-thalamo-cortical loop; EPI = echo planar imaging; ER = event related; FFA = flexible factorial analysis; FIR = finite impulse response; fMRI = functional magnetic resonance imaging; FWE = family-wise error; FWHM = full width at half maximum; GLM = general linear model; GP = globus pallidus; HRF = hemodynamic response function; M1 = primary motor cortex; MNI = Montreal Neurological Institute; MP-RAGE = Magnetization-Prepared RApid Gradient Echo; $n$ = number of cases; $p$ = error probability; PD = Parkinson's disease; PMC = premotor cortex; Pu = putamen; ROI = region of interest; S1 = primary somatosensory cortex; SMA = supplementary motor area; SN = substantia nigra; $T_1$ = longitudinal relaxation time; Thal = thalamus; $TR$ = repetition time; UPDRS = unified Parkinson's disease rating scale.

### Supplementary Tables

**Supplementary Table S1 |** One-sample *t*-test results (MNI coordinates in mm and *t*-values of the peak activation in a cluster as well as cluster sizes and FWE-corrected *p*-values) for finger tapping with the (non-dominant) **left hand** obtained in the initial (*n*=11), new (*n*=20) and combined cohort (*n*=31) with the standard model (block design employing a boxcar function), the glove model, and the onset model. Corresponding activation maps are shown in Supplementary Figure S2.

| Region | Standard model | | | | | | Glove model | | | | | | Event-related model | | | | | |
|---|---|---|---|---|---|---|---|---|---|---|---|---|---|---|---|---|---|---|
| | Peak | | | | Cluster | | Peak | | | | Cluster | | Peak | | | | Cluster | |
| | *x* | *y* | *z* | *t*-val. | $N_{vox}$ | $p_{FWE}$ | *x* | *y* | *z* | *t*-val. | $N_{vox}$ | $p_{FWE}$ | *x* | *y* | *z* | *t*-val. | $N_{vox}$ | $p_{FWE}$ |
| *Initial cohort (n = 11)* | | | | | | | | | | | | | | | | | | |
| Right M1 | | | | | | | 46 | –16 | 56 | 11.39 | 138 | 0.006 | | | | | | |
| *New cohort (n = 20)* | | | | | | | | | | | | | | | | | | |
| Right M1 | 48 | –16 | 56 | 8.19 | 45 | <0.001 | 36 | –20 | 64 | 7.45 | 210 | 0.001 | | | | | | |
| Right PM/SMA | 34 | –22 | 68 | 5.70 | | <0.010 | | | | | | | | | | | | |
| Right S1 | | | | | | | 54 | –16 | 52 | 5.63 | | 0.011 | 52 | –18 | 58 | 6.63 | 48 | 0.001 |
| *Combined cohort (n = 31)* | | | | | | | | | | | | | | | | | | |
| Right M1/PM/SMA | 48 | –16 | 56 | 10.09 | 186 | <0.001 | 40 | –16 | 58 | 10.47 | 684 | <0.001 | 36 | –22 | 68 | 6.08 | 153 | 0.010 |
| Right S1 | | | | | | | | | | | | | 52 | –18 | 58 | 8.70 | | <0.001 |

Abbreviations: M1 = primary motor cortex; PM = premotor cortex; S1 = primary somatosensory cortex; SMA = supplementary motor area.

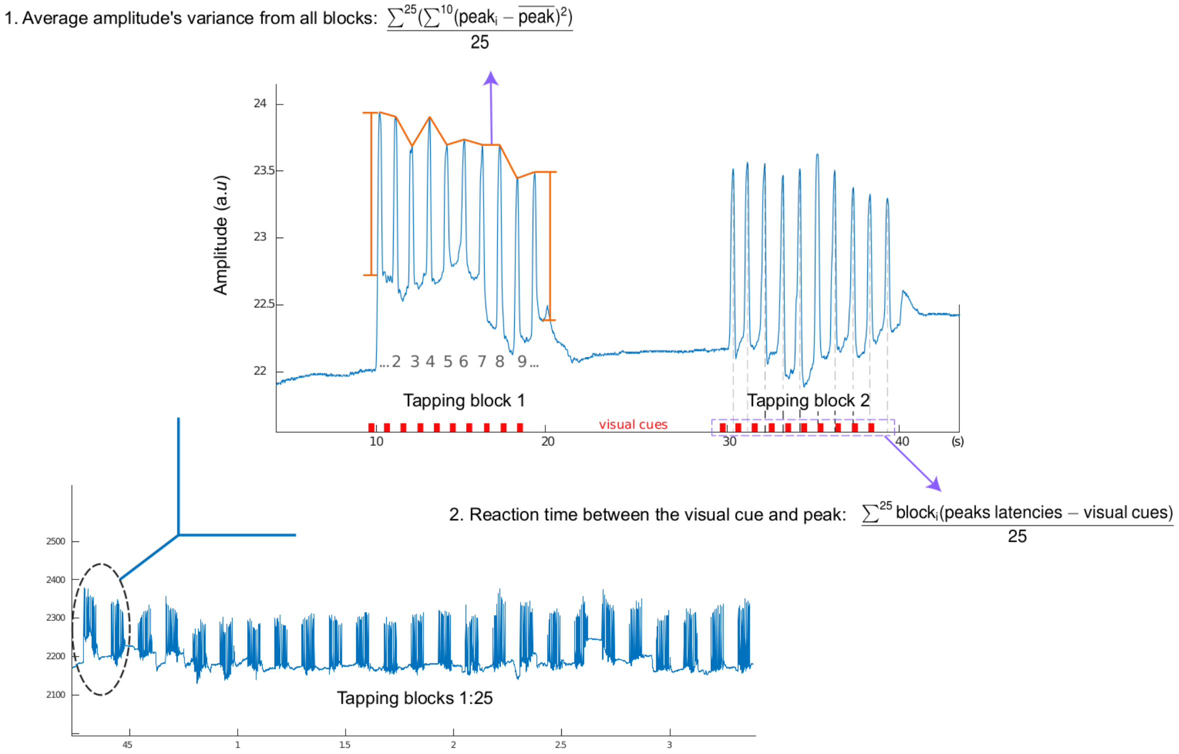


**Supplementary Figure S1 |** Depiction of the two parameters characterizing finger tapping, which were used in the analysis of the behavioral data.

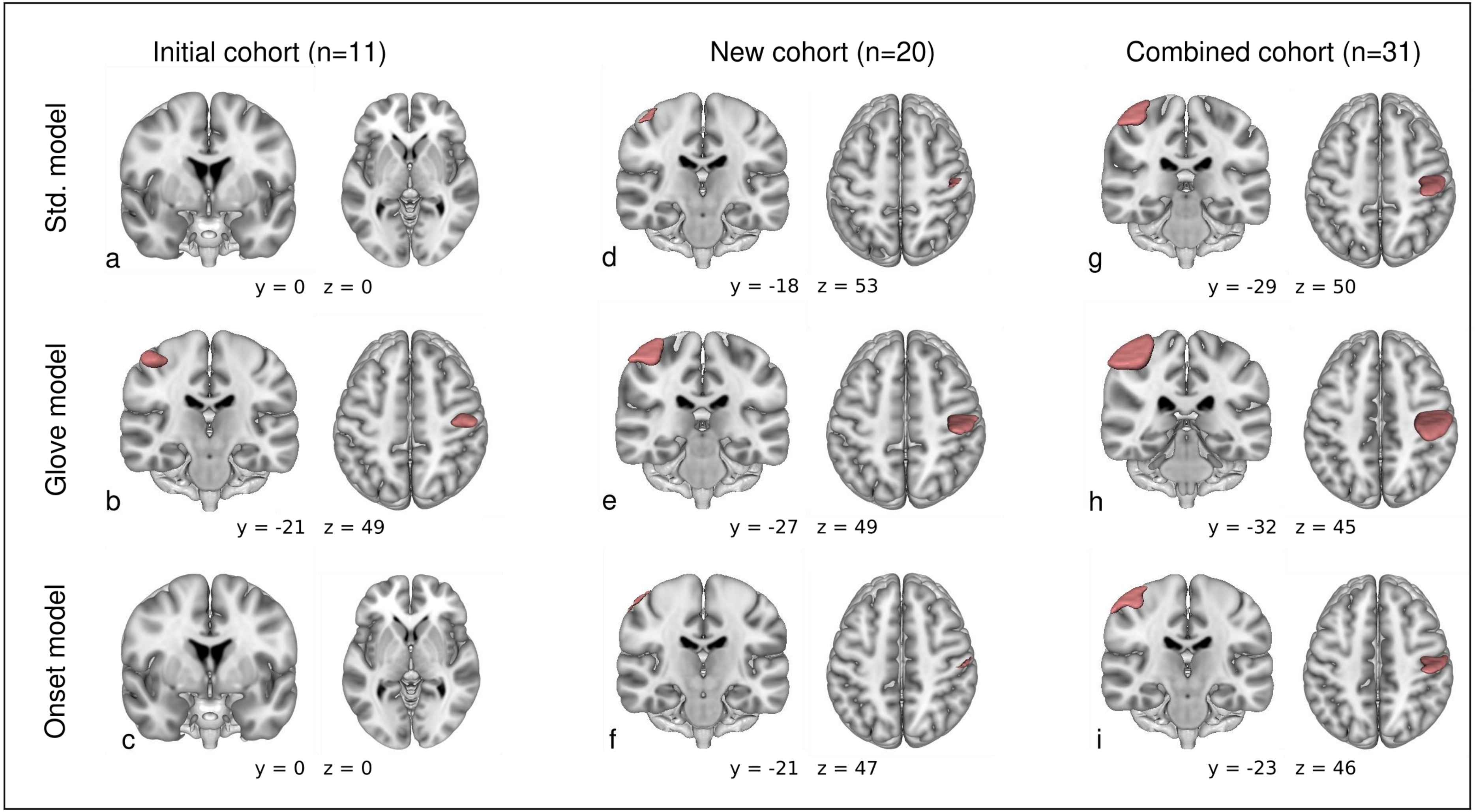


**Supplementary Figure S2 |** Activation maps (main effect of tapping vs. rest; group results) for finger tapping with the (non-dominant) **left hand** (L-DOPA ON) obtained in the initial (*n*=11) **(a–c)**, new (*n*=20) **(d–f)** and combined cohort (*n*=31) **(g–i)** with the standard model (block design employing a boxcar function) **(a, d, g)**, the glove model **(b, e, h)**, and the onset model **(c, f, i)**. The coordinates refer to displayed anatomical slices and not to the clusters' maxima. Further quantitative results are summarized in Supplementary Table S1.
